# Elucidating Tumor-stromal Metabolic Crosstalk in Colorectal Cancer through Integration of Constraint-Based Models and LC-MS Metabolomics

**DOI:** 10.1101/2021.05.28.446227

**Authors:** Junmin Wang, Alireza Delfarah, Patrick Gelbach, Emma Fong, Paul Macklin, Shannon M. Mumenthaler, Nicholas A. Graham, Stacey Finley

## Abstract

Colorectal cancer (CRC) is a major cause of morbidity and mortality in the United States. Tumor-stromal metabolic crosstalk in the tumor microenvironment promotes CRC development and progression, but exactly how stromal cells, in particular cancer-associated fibroblasts (CAFs), affect the metabolism of tumor cells remains unknown. Here we take a data-driven approach to investigate the metabolic interactions between CRC cells and CAFs, integrating constraint-based modeling and metabolomic profiling. Using metabolomics data, we perform unsteady-state parsimonious flux balance analysis to infer flux distributions for central carbon metabolism in CRC cells treated with or without CAF-conditioned media. We find that CAFs reprogram CRC metabolism through stimulation of glycolysis, the oxidative arm of the pentose phosphate pathway (PPP), and glutaminolysis as well as inhibition of the tricarboxylic acid cycle. To identify potential therapeutic targets, we simulate enzyme knockouts and find that inhibiting the hexokinase and glucose-6-phosphate dehydrogenase reactions exploits the CAF-induced dependence of CRC cells on glycolysis and oxidative PPP. Our work gives mechanistic insights into the metabolic interactions between CRC cells and CAFs and provides a framework for testing hypotheses towards CRC-targeted therapies.

## 1. Introduction

Colorectal cancer (CRC) remains one of the deadliest cancers in the United States, with a 5-year survival rate of less than 15% for patients with stage IV CRC [1]. Over 140,000 people are diagnosed with CRC each year, leading to approximately 50,000 deaths [2]. Interactions between tumor and stromal cells have long been speculated to promote tumor development and progression. Cancer-associated fibroblasts (CAFs), a dominant cellular component of the tumor stroma, play a significant role in cancer pathogenesis by contributing to the cancer cells’ altered metabolism, a hallmark of CRC [3, 4]. Various factors secreted by CAFs, including hepatocyte growth factor (HGF) and neuregulin-1, are known to inhibit therapeutic response in cancer [5, 6]. In addition, increased deposition of matrix proteins (e.g., hyaluronan and collagen) by CAFs has been found to affect drug penetration [7, 8, 9].

Increasing evidence supports the idea of reciprocal metabolic reprogramming among CRC cells and CAFs, but questions remain regarding the mechanism of the metabolic dependencies. For example, it is not clear if the influence of CAFs causes CRC cells to redistribute their carbon fluxes through central carbon metabolism, and whether oncogenes such as KRAS contribute to the metabolic reprogramming. Understanding the characteristics of tumor cells and CAFs in their metabolic ecosystem may provide insight needed to develop optimal cancer therapies.

Here, we applied a systems biology approach that combines metabolomics with constraint-based models to understand how CAFs influence CRC cell metabolism. We profiled metabolic alterations in KRAS^WT^ and KRAS^MUT^ DLD-1 cells (a CRC cell line) either cultured solely in CRC media or cultured initially in CRC media and then switched to CAF-conditioned media (Figure 1). Metabolite pool size profiling alone, however, does not explain how cells modulate their reaction fluxes to achieve the balance between anabolism and catabolism. To infer flux distributions, we developed unsteady-state parsimonious flux balance analysis (upFBA), a data-driven modeling approach that integrates constraint-based methods and liquid chromatography-mass spectrometry (LC-MS) metabolomics data [10, 11]. We constructed a network of central carbon metabolism similar to [12] and performed upFBA to estimate the intracellular reaction fluxes of KRAS^WT^ and KRAS^MUT^ CRC cells grown in CRC media and CAF-conditioned media.

**Figure 1:**
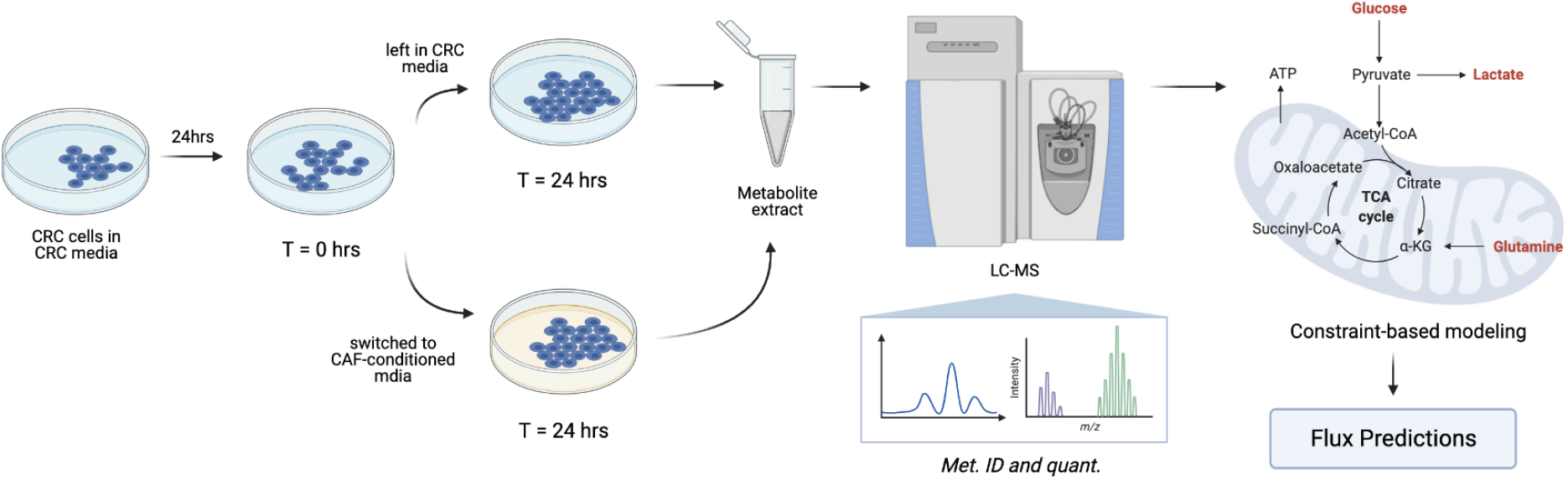
Experimental workflow of our study. KRAS^WT^ and KRAS^MUT^ CRC cells were first cultured in CRC media (DMEM) for 24 hrs. At T = 0 hrs, they were either left in their normal media, i.e., CRC media or switched to CAF-conditioned media for another 24 hrs. At T = 24 hrs, metabolites were extracted from cells and from the media and subjected to LC-MS analysis. For all conditions (KRAS^WT^ CRC cells in CRC media, KRAS^MUT^ CRC cells in CRC media, KRAS^WT^ CRC cells in CAF-conditioned media, and KRAS^MUT^ CRC cells in CAF-conditioned media), we measured the fold changes of intracellular metabolites and the secretion/uptake rates of glucose, lactate, and glutamine. These measurements were used to constrain an upFBA model we developed to infer the flux distributions of central carbon metabolism in CRC cells. (See Section 4 for details). This figure was created from BioRender.com

Our analysis indicates that CAFs play a pivotal role in regulating CRC metabolism through stimulation of glycolysis, the oxidative arm of the pentose phosphate pathway (PPP), and glutaminolysis, as well as inhibition of the tricarboxylic acid (TCA) cycle. Concomitantly, CAFs induce a higher flux through substrate-level phosphorylation and NADH production. To identify potential therapeutic targets, we subsequently performed unsteady-state flux balance analysis (uFBA) in search of gene deletions that could lead to reduced cancer growth. In particular, we found that inhibition of the hexokinase (HK) or glucose-6-phosphate dehydrogenase (G6PD) reactions results in reduced cancer cell growth in CAF-conditioned media compared to CRC media by exploiting the CRC-CAF metabolic crosstalk. Our results suggest that CAFs make CRC cells more dependent on glycolysis and PPP and hence more vulnerable to the inhibition of key regulatory enzymes of those metabolic pathways.

## 2. Results

### 2.1. LC-MS Profiling Reveals Media-Induced Metabolic Alterations

To directly assess changes in the levels of CRC cells’ intracellular metabolites, we performed a semi-targeted LC-MS metabolomics analysis of central carbon metabolism in KRAS^WT^ and KRAS^MUT^ CRC cells cultured in CRC media or CAF-conditioned media. (See Sections 4.1 and 4.2 for details). Altogether, 81 metabolites were consistently identified and quantified across all samples.

We calculated the fold changes of the metabolite levels and applied moderated *t*-tests to validate the differential abundance of metabolites [13]. (See Section 4.2 for details). A metabolite was considered significantly different between two groups if its fold change was larger than 1.2 and false discovery rate (FDR) was less than 0.05. A total of 18 metabolites were considered to be significantly different between CRC media and CAF-conditioned media for KRAS^WT^ CRC cells; a total of 20 metabolites differed significantly for KRAS^MUT^ CRC cells (Figure 2(a), (b)). In contrast, only 5 metabolites were significantly different between KRAS^WT^ and KRAS^MUT^ cells in CRC media; only 2 metabolites differed significantly in CAF-conditioned media (Figure 2(c), (d)). This suggests that CAFs induce more pronounced alterations in CRC metabolism than oncogenic KRAS, as cells of different genotype identities show fewer differences in their metabolic profiles compared to cells grown in different media (Figure 2(a)-(d)).

**Figure 2:**
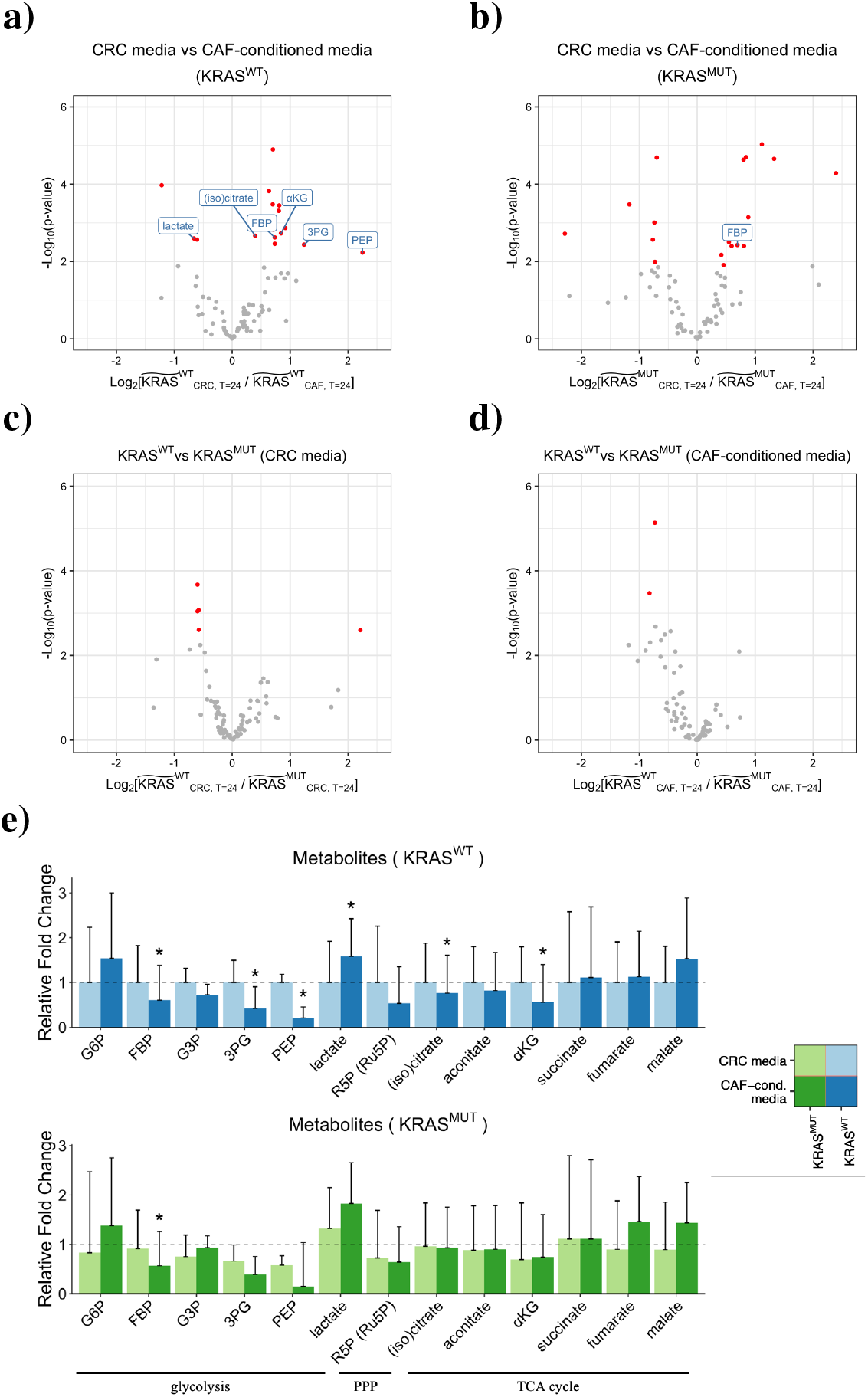
Distinct metabolic profiles of CRC cells grown under different conditions. a) - d) Comparison of intracellular metabolites between two conditions. The wide tilde in the *x*-axis label represents the fold change in metabolite abundance between T = 0 hrs and T = 24 hrs. Metabolites that are significantly different between two groups are highlighted in red (FDR *<* 0.05, fold change *>* 1.2). Metabolites unique to glycolysis, PPP, and the TCA cycle are labeled. e) Fold changes of metabolites belonging to glycolysis, PPP, and the TCA cycle in KRAS^WT^ (top) and KRAS^MUT^ (bottom) CRC cells. (Glc, 6PG, and OAA are not shown due to high coefficients of variation). The heights of the bars represent mean fold changes, and error bars represent standard deviations. A metabolite is considered significantly different between CRC media and CAF-conditioned media if its fold change is larger than 1.2 and FDR is less than 0.05 (asterisks above bars).

Grouping metabolites by pathways (i.e., glycolysis, PPP, and the TCA cycle), we found that no sets of metabolites unique to a particular pathway were consistently regulated in one direction, either up or down (Figure 2(e)). This, however, does not necessarily indicate that CAFs do not influence CRC metabolism. Metabolic networks are highly intertwined, and for most metabolites, synthesis and breakdown take place simultaneously. The accumulation of a pathway intermediate can be caused by upregulated anabolism and/or downregulated catabolism. Thus, we sought to look at pathway fluxes, rather than individual metabolite levels.

### 2.2. A Data-Driven Model Predicts Metabolic Fluxes in CRC

To infer intracellular flux distributions, we used a constraint-based model of central carbon metabolism in CRC cells. Our model was adapted from the constraint-based model developed by [12] and included the same 89 metabolites and 73 reactions as described in [12], including the biomass template reaction. In light of glutamine addiction that CRC cells are shown to exhibit [14], we included an additional reaction to account for the conversion of glutamate to glutamine mediated by glutamine synthetase (GS) [15]. We performed random sampling of mass balance constraints based on knowledge of initial metabolite concentrations from literature and measurements of fold changes of metabolite concentrations between T = 0 hrs and T = 24 hrs from experimental data. (See Section 4.3 for details and Table S1 for initial metabolite concentrations). Biomass growth rates, which we calculated by fitting an exponential growth curve to cell count measurements taken over 3 days, were used to constrain our models. (See Sections 4.1 and 4.3 for details and Table S2 for data). The experimentally measured extracellular uptake/secretion rates of glucose, lactate, and glutamine were also used to constrain our models. (See Sections 4.2 and 4.3 for details and Table S2 for data).

For each set of mass balance constraints, we performed upFBA in search of a flux distribution that minimizes the total sum of fluxes through the network [10, 11]. Based on the calculated fluxes, we separated reactions into essential reactions and non-essential reactions. Essential reactions are defined as reactions that maintain a non-zero flux under at least one set of mass balance constraints, and non-essential reactions are those that do not. The resulting minimized total sums of fluxes, numbers of essential reactions, and numbers of non-essential reactions are given in Figure S1. We did not find significant differences in the total flux through the metabolic reactions or the number of essential reactions across the four conditions. Differential flux was validated on all essential reactions using the nonparametric Wilcoxon rank sum test [16]. (Details can be found in Section 4.3). We visualized the distributions of the median fluxes via Voronoi treemaps in Figure S2. It can be seen that under all conditions, glycolysis accounts for the largest proportion of the total sum of fluxes among the pathways (Figure S2). Median fluxes, median absolute deviations (MADs), and adjusted p-values are provided in Supplementary File a. Figures 3, 4, and 5 summarize the predicted flux distributions of CRC cells grown in CRC media and CAF-conditioned media. We next discuss in detail these predicted fluxes for glycolysis, PPP, TCA cycle, and glutaminolysis reactions and compare the flux values between the four experimental conditions.

**Figure 3:**
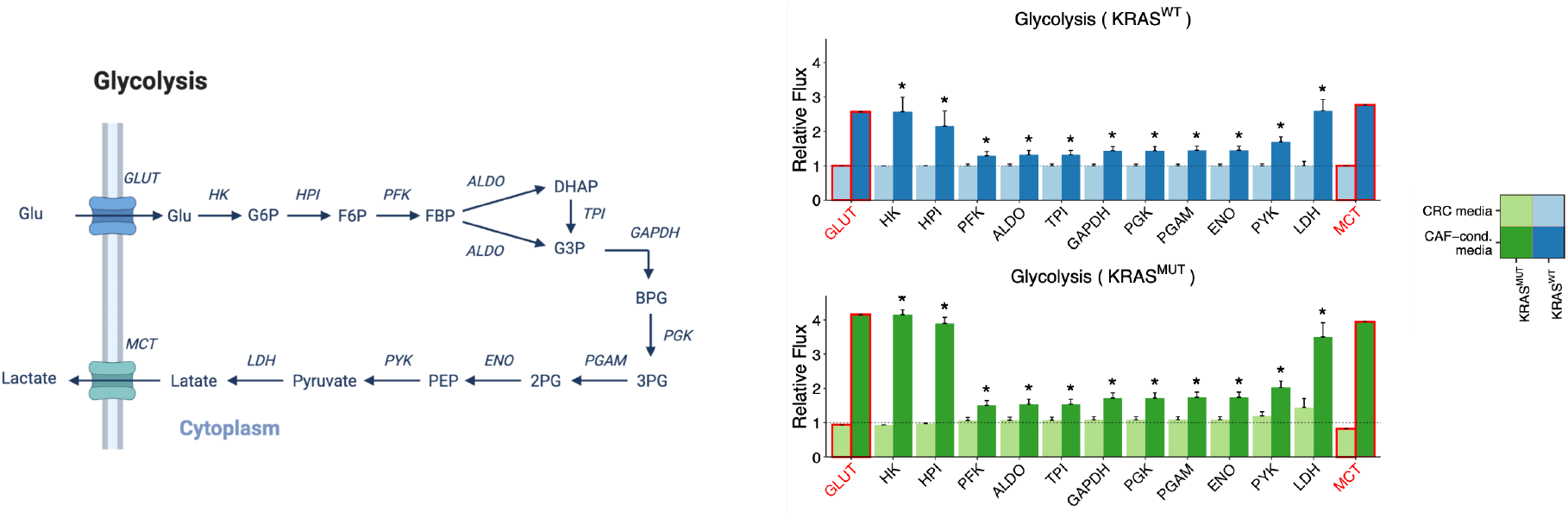
Relative fluxes of glycolysis in KRAS^WT^ and KRAS^MUT^ CRC cells cultured in CRC media compared to CAF-conditioned media. *Left*, Schematic of the glycolysis pathway for reference. This schematic was created from BioRender.com. Directions of the arrows in the diagram represent directions of the reactions. *Right*, Predicted fluxes for glycolysis reactions. All reactions have been normalized to the respective flux in KRAS^WT^ CRC cells cultured in CRC media. The heights of the bars represent normalized median fluxes, and error bars represent normalized MADs. A flux is considered significantly different between CRC media and CAF-conditioned media if its fold change is larger than 1.2 and adjusted p-value is less than 0.001 (asterisks above bars). (We lower the p-value threshold in order to account for the larger sample size we achieved via simulations than experiments.) We outline in red the fluxes through GLUT and MCT, which are experimentally measured and supplied to the model as fixed constraints.

**Figure 4:**
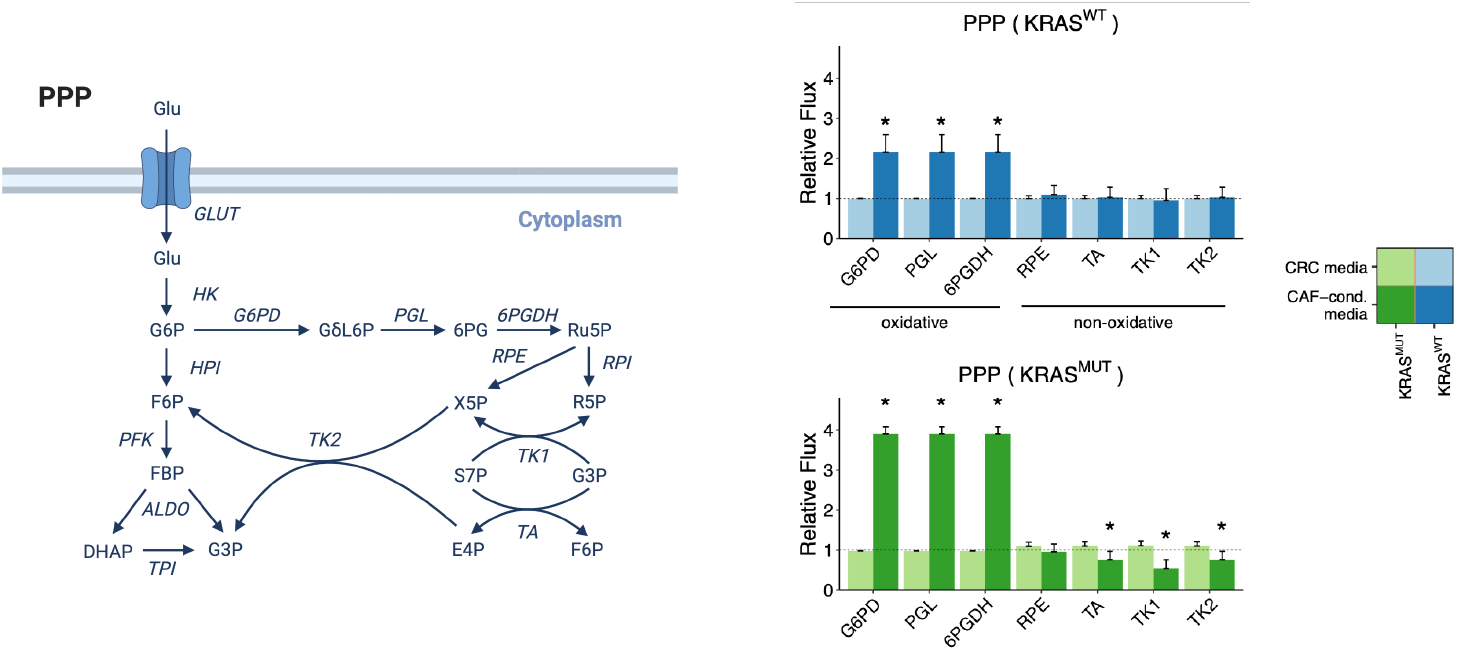
Relative fluxes of PPP in KRAS^WT^ and KRAS^MUT^ CRC cells cultured in CRC media compared to CAF-conditioned media. *Left*, Schematic of the PPP for reference. This schematic was created from BioRender.com. Directions of the arrows in the diagram represent directions of the reactions. *Right*, Predicted fluxes for PPP reactions. All reactions have been normalized to the respective flux in KRAS^WT^ CRC cells cultured in CRC media. The heights of the bars represent normalized median fluxes, and error bars represent normalized MADs. A flux is considered significantly different between CRC media and CAF-conditioned media if its fold change is larger than 1.2 and adjusted p-value is less than 0.001 (asterisks above bars).

**Figure 5:**
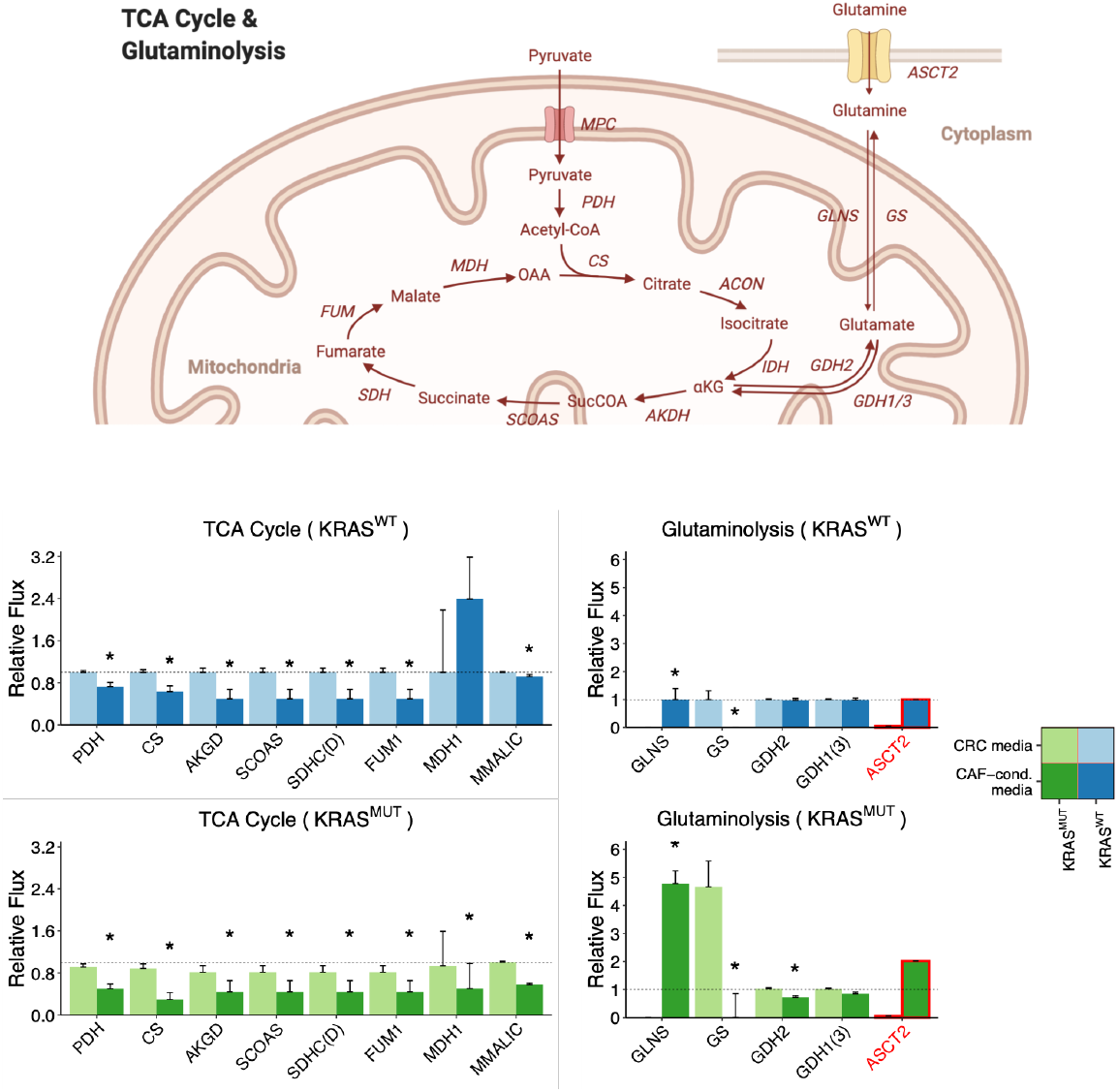
Relative fluxes of the TCA cycle and glutaminolysis. KRAS^WT^ and KRAS^MUT^ CRC cells cultured in CRC media are compared to CAF-conditioned media. *Top*, Schematic of the TCA cycle and glutaminolysis pathways for reference. This schematic was created from BioRender.com. Directions of the arrows in the diagram represent directions of the reactions. *Bottom*, Predicted fluxes for TCA cycle and glutaminolysis reactions. All reactions except GLNS and ASCT2 have been normalized to the respective flux in KRAS^WT^ CRC cells cultured in CRC media. For visualization, GLNS and ASCT2 have been normalized to the respective flux in KRAS^WT^ cells cultured in CAF-conditioned media. The heights of the bars represent normalized median fluxes, and error bars represent normalized MADs. A flux is considered significantly different between CRC media and CAF-conditioned media if its fold change is larger than 1.2 and adjusted p-value is less than 0.001 (asterisks above bars). We outline in red the flux through ASCT2, which is experimentally measured and supplied to the model as a fixed constraint.

### 2.3. CAFs Enhance Glycolytic Fluxes of CRC Cells

Our extracellular metabolomics data indicate that KRAS^WT^ and KRAS^MUT^ CRC cells cultured in CRC media have comparable glucose uptake and lactate secretion rates, which are mediated by GLUT and MCT, respectively in our model (Figure 3). (See Table S2 for data). When switched to CAF-conditioned media, KRAS^WT^ and KRAS^MUT^ cells increase glucose uptake rates by 2.57- and 4.41-fold, respectively, and lactate secretion rates increase by 2.77- and 4.77-fold. This indicates that CAFs play a stimulatory role in glucose uptake and lactate secretion. Similar to CAFs, oncogenic KRAS is reported to upregulate glucose uptake and lactate secretion [17, 18]. This effect, however, is only pronounced in CAF-conditioned media, as the glucose uptake and lactate secretion rates differ to a much lesser extent between KRAS^WT^ and KRAS^MUT^ cultured in the CRC media (Figure 3).

Regarding intracellular fluxes, upFBA predicts that switching to CAF-conditioned media significantly increases the fluxes through all the glycolytic reactions in both KRAS^WT^ and KRAS^MUT^ CRC cells (Figure 3). This suggests that CAFs can upregulate the glycolytic fluxes of CRC cells, and that this interaction occurs by secreted factors and not by contact. Although glucose uptake rates are comparable in KRAS^WT^ and KRAS^MUT^ cells cultured in CRC media, the flux through the lactate dehydrogenase (LDH) reaction, which dictates lactate secretion, increases by 1.45-, 2.60-, and 3.50-fold, respectively in KRAS^MUT^ cells cultured in CRC media, KRAS^WT^ cells grown in CAF-conditioned media, and KRAS^MUT^ cells cultured in CAF-conditioned media compared to KRAS^WT^ cells grown in CRC media (Figure 3). This suggests that both oncogenic KRAS and CAFs can upregulate lactate fermentation, although CAFs show a stronger effect.

### 2.4 CAFs Promote NADPH Production in CRC Cells through Oxidative PPP

The PPP maintains carbon homeostasis by synthesizing the ribose ring of nucleotides and providing reducing equivalents in the form of NADPH [19, 20]. upFBA predicts that KRAS^WT^ and KRAS^MUT^ CRC cells cultured in CAF-conditioned media have higher fluxes through reactions unique to the oxidative arm of the PPP, i.e., G6PD, lactonase (PGL), as well as 6-phosphogluconate dehydrogenase (6PGDH), than their counterparts cultured in CRC media (Figure 4). In contrast, the non-oxidative arm of the PPP, comprised of ribose phosphate epimerase (RPE), transaldolase (TA), and transketolases (TK1 and TK2), is barely affected by the growth media in KRAS^WT^ cells (Figure 4). Note that, TK1 proceeds in the direction of S7P → R5P (Figure 4), which indicates that non-oxidative is also directed to support R5P production in CRC cells. Reduced flux through the TA, TK1, and TK2 reactions in KRAS^MUT^ cells in CAF-conditioned media suggests that CRC cells can decouple the oxidative and non-oxidative arms of the PPP to maintain precise control of NAPDH and R5P levels. A similar phenomenon has been observed previously in pancreatic cancer cells [18]. This decoupling is not observed in KRAS^WT^ cells, revealing a unique contribution of the KRAS mutation to CRC cell metabolism.

### 2.5. CAFs Inhibit the TCA Cycle and Glutamine Anabolism of CRC Cells

In eukaryotic cells, pyruvate produced at the end of glycolysis can be transported into the mitochondria to fuel the TCA cycle. Unlike glycolysis and oxidative PPP, we found that switching to CAF-conditioned media inhibits the TCA cycle in CRC cells, as the fluxes through the TCA cycle are in general predicted to be lower for cells in CAF-conditioned media, compared to CRC media (Figure 5).

In addition to being an important biosynthetic pathway, the TCA cycle also serves to connect glucose and amino acid metabolism. Glutamate, which can be synthesized from glutamine, enters the TCA cycle through its conversion to *α*-ketoglutarate via glutamate dehydrogenase (GDH) [21]. We observed a distinct difference in how CRC cells utilize glutamine in CRC media and CAF-conditioned media, as the measured glutamine uptake rates of KRAS^WT^ and KRAS^MUT^ CRC cells cultured in CAF-conditioned media are 16 and 39 times higher, respectively, than the glutamine uptake rates of their counterparts cultured in CRC media (Figure 5). (See Table S2 for data. Glutamine uptake is mediated by ASCT2 in our model). Furthermore, upFBA predicts that the increased glutamine uptake rates induced by CAFs lead to reprogrammed intracellular glutamine metabolism. CRC cells cultured in CAF-conditioned media divert excessive glutamine via amidohydrolysis mediated by glutaminase (GLNS), whereas CRC cells cultured in CRC media compensate for the reduced glutamine uptake by elevating the flux through the reaction converting glutamate to glutamine, which is mediated by the GS enzyme (Figure 5). Note that enhanced glutaminolysis does not result in elevated fluxes through the TCA cycle (Figure 5). This suggests that CAF-induced increases in glutamine uptake rates are perhaps not driven by the TCA cycle anaplerosis, but by a rising demand for glutamate needed to fulfill other cellular functions, such as transamination and glutathione synthesis [22].

### 2.6. CAFs Induce Reprogramming of NADH, ATP, and Pyruvate Metabolism in CRC Cells

To understand how CAF-induced metabolic alterations affect energy production, we quantified how individual reactions contribute to the production of NADH and ATP as well as the consumption of pyruvate under different conditions. Upon hydrogen transfer to oxygen in oxidative phosphorylation, each NADH maximally produces three ATP molecules, so electron transport from NADH produced in the TCA cycle is widely considered a major source of cellular ATP [23]. upFBA predicts a stimulatory role of CAFs in NADH production, as the total flux of NADH production is higher in CRC cells cultured in CAF-conditioned media than those cultured in CRC media (Figure 6(a)). Among the contributing enzymes, glyceraldehyde-3-phosphate dehydrogenase (GAPDH) is predicted to be the most active, and flux through this reaction further increases in the presence of CAFs (Figure 6(a)). As is shown in Figure 6(a), GAPDH not only has a higher absolute flux, but also accounts for a larger fraction of NADH production in CAF-conditioned media.

**Figure 6:**
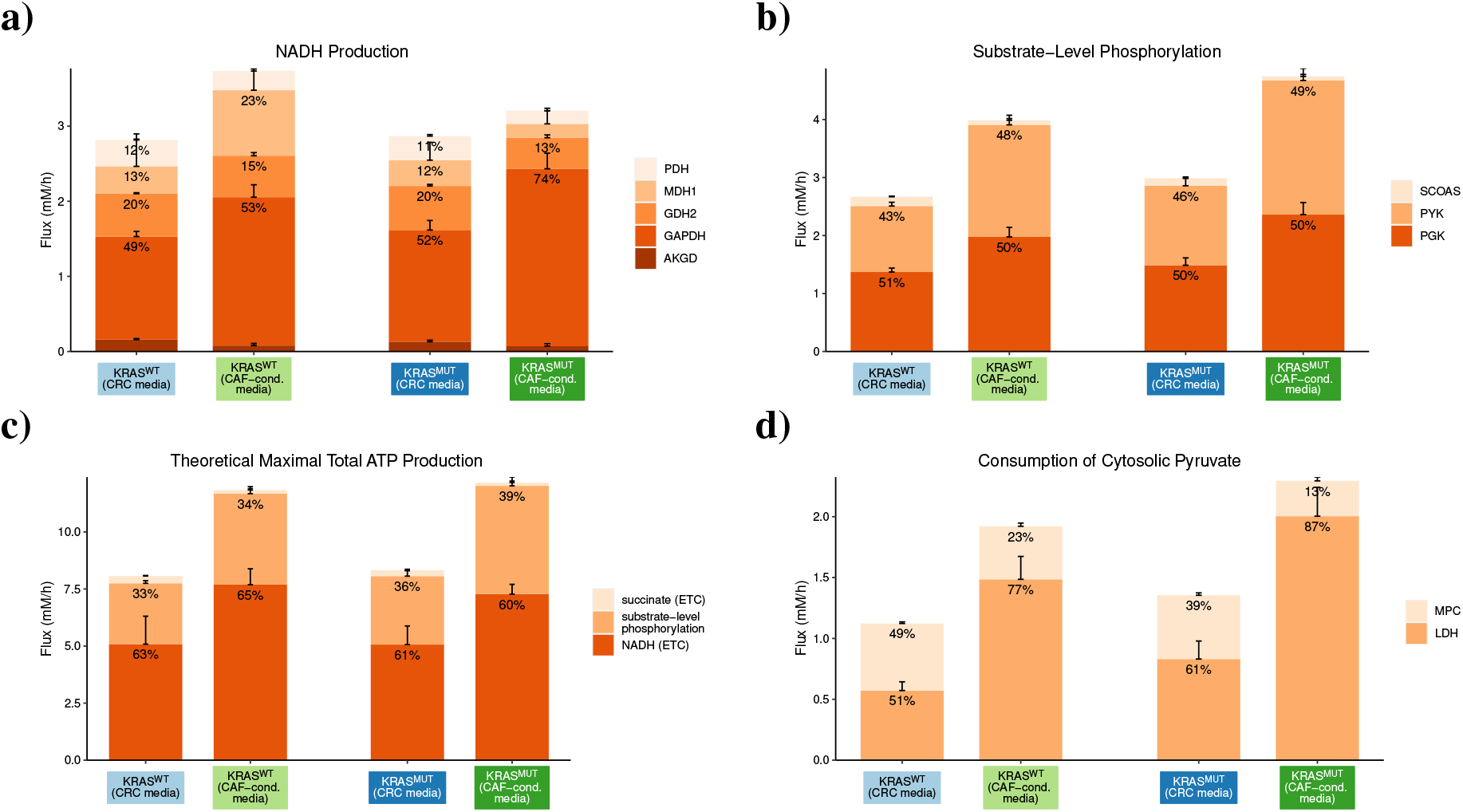
Predicted flux splits of metabolite species. a) NADH production; b) substrate-level phosphorylation; c) theoretical maximum yield of ATP; and d) cytosolic pyruvate consumption in CRC cells. The heights of the bars represent the total sums of absolute flux values. Colors represent enzyme-mediated reactions. Numbers at the top of each stacked bar represent the relative fractions of the total flux devoted to a reaction (*<* 8% not shown). Error bars on top of each stacked bar represent the MADs of the fluxes.

Besides oxidative phosphorylation, another source of cellular ATP is substrate-level phosphorylation. Although the relative contributions of the succinyl-CoA synthetase (SCOAS), pyruvate kinase (PYK), and phosphoglycerate kinase (PGK) reactions to overall substrate-level phosphorylation are very similar across all four conditions, the absolute fluxes through these reactions are higher in KRAS^MUT^ CRC cells and CAF-conditioned media than in KRAS^WT^ CRC cells and CRC media (Figure 6(b)). Assuming that oxidation of one molecule of NADH or succinate through the electron transport chain (ETC) produces three or two molecules of ATP, respectively, [23], we then calculated the theoretical total maximum yield of ATP from the given fluxes of reactions that produce ATP and ATP precursors in central carbon metabolism. As is shown in Figure 6(c), the theoretical total maximum flux of ATP production is higher in CAF-conditioned media than in CRC media. The enhanced theoretical total maximum production of ATP in CAF-conditioned media suggests that CAFs may induce CRC cells to raise the level of ATP production through both substrate-level phosphorylation and oxidative phosphorylation. Note that a higher theoretical total maximum flux of ATP production does not necessarily indicate a higher ATP level, as ATP level is determined by both ATP production and consumption.

Though not a direct energy-carrying molecule, pyruvate serves as a key branching point of central metabolism: it can either be converted to lactate via LDH or transported into the mitochondria via mitochondrial pyruvate carriers (MCPs). upFBA predicts that both oncogenic KRAS and CAFs can enhance lactate fermentation, as the flux through LDH-mediated conversion, both in terms of absolute values and relative fractions, is higher in KRAS^MUT^ cells and CAF-conditioned media than in KRAS^WT^ cells and CRC media (Figure 6(d)).

### 2.7. Gene Deletion Analysis Predicts Knockout Metabolic Phenotypes

Next, we applied flux models to examine how the cancer cells’ growth rate changes in response to enzyme knockouts. To simulate a single-enzyme knockout, we constrained the flux of one reaction in the original metabolic phenotype to zero. Subsequently, we performed uFBA in search of flux distributions that maximize the biomass growth rates under the new constraints. (See Section 4.3 for details). Given that there are 100 sets of mass balance constraints, 74 reactions, 2 genotype identities (KRAS^WT^ and KRAS^MUT^), and 2 media conditions (CRC media and CAF-conditioned media), we simulated a total of 100 × 74 × 2 × 2 = 29600 single-enzyme knockouts.

We compared the predictions of the median maximal biomass growth rates upon gene deletion between CRC media and CAF-conditioned media. Similar to the comparison of original metabolic phenotypes, we validated the differences in biomass growth rates via the Wilcoxon rank sum test [16]. (See Section 4.3 for details). Median maximal biomass growth rates of single-enzyme-knockout phenotypes, MADs, and adjusted p-values are provided in Supplementary File b. Among a total of 25 enzyme knockouts leading to different biomass growth rates in KRAS^WT^ CRC cells, 16 belong to central carbon metabolism: 9, 5, and 2 are unique to glycolysis, PPP, and the TCA cycle, respectively (Figure 7). Similarly, out of 19 enzyme knockouts that lead to different biomass growth rates in KRAS^MUT^ CRC cells, 15 belong to central carbon metabolism, with 9, 4, and 2 enzyme knockouts from glycolysis, PPP, and the TCA cycle, respectively. In both KRAS^WT^ and KRAS^MUT^ CRC cells, 11 single-enzyme-knockouts belonging to central carbon metabolism result in reduced biomass growth rates in CRC media. In contrast, only 5 single-enzyme-knockouts result in reduced biomass growth rates in CAF-conditioned media for KRAS^WT^ CRC cells, and only 4 single-enzyme-knockouts, for KRAS^MUT^ CRC cells (Figure 7). As CAFs are generally believed to make cancer cells more resilient to enzyme knockouts, it is unsurprising that fewer enzyme knockouts are available for blocking cancer growth in CAF-conditioned media. Nevertheless, it is noteworthy that for both KRAS^WT^ and KRAS^MUT^ cells, knockouts of HK and G6PD, traditionally considered the rate-limiting steps of glycolysis and PPP, result in lower biomass growth rates in CAF-conditioned media compared to CRC media (Figure 7). This is likely because CAFs make CRC cells more dependent on glycolysis and PPP, as we have shown in Sections 2.3 and 2.4. Thus, interference with glycolysis and PPP could take a heavier toll on CRC metabolism in presence of CAFs.

**Figure 7:**
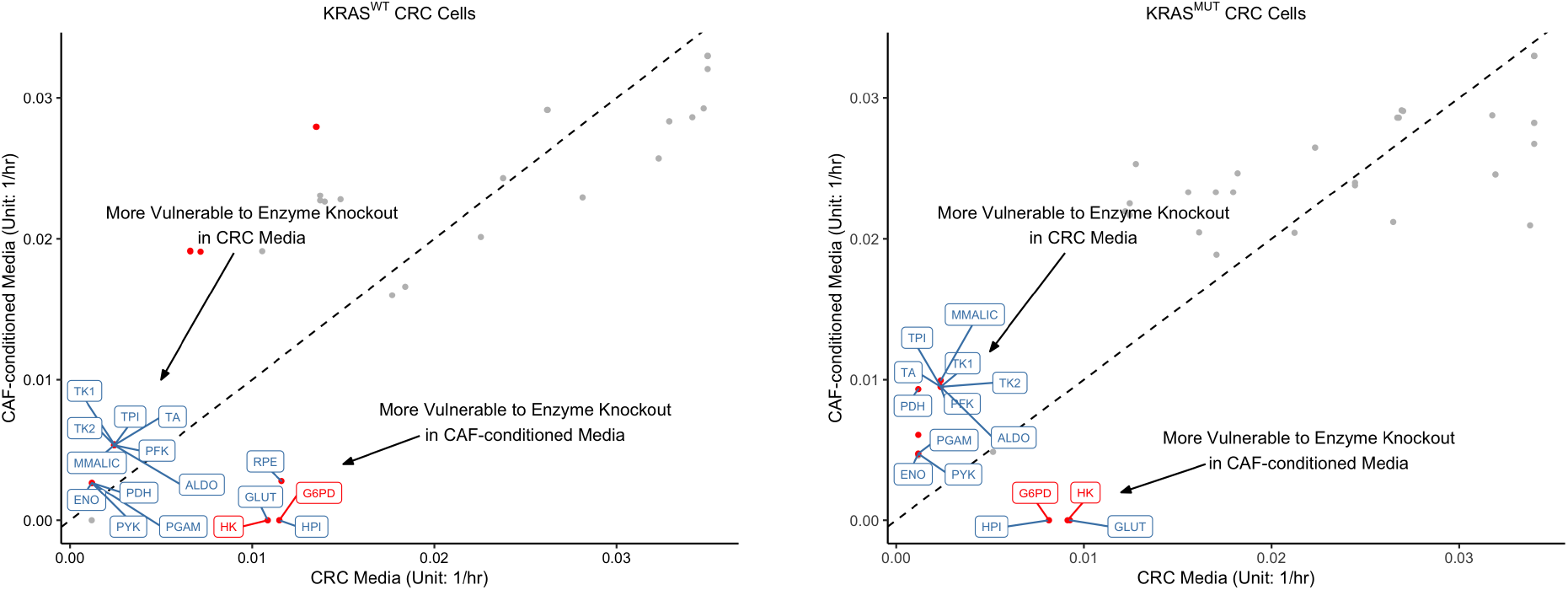
Predicted maximal biomass growth rates upon gene deletions. uFBA was used to predict the maximal growth rates for KRAS^WT^ and KRAS^MUT^ CRC cells upon gene deletions. The *x*- and *y*-coordinates of each point represent the median maximal biomass growth rates of CRC cells cultured in CRC media and CAF-conditioned media, respectively when the same enzyme is inhibited. The dashed diagonal line represents equal median rates in both media. Gene deletions resulting in significantly different biomass growth rates between CRC media and CAF-conditioned media are represented by red dots (adjusted p-value *<* 0.001, fold change *>* 2). Enzymes belonging to glycolysis, PPP, the TCA cycle, and glutaminolysis are labeled.

In contrast to the effects of the medium, few single-enzyme knockouts lead to different biomass growth rates between KRAS^WT^ and KRAS^MUT^ cells (Figure S3). In CRC media, the inhibition of RPE is the only enzyme knockout that results in differential cancer growth rates between KRAS^WT^ and KRAS^MUT^ CRC cells (reduced growth in KRAS^MUT^ cells; Figure S3). In CAF-conditioned media, only three enzyme knockouts, including the knockouts of GAPDH and pyruvate dehydrogenase (PDH), lead to differential cancer growth rates between the two cell lines (Figure S3). (All lead to reduced growth in KRAS^WT^ cells). This is consistent with our findings in Sections 2.1 - 2.5 and further shows that CAF-conditioned media cause stronger metabolic alterations to CRC cells than oncogenic KRAS.

## 3. Discussion

In this work, we have studied how CAFs affect CRC metabolism via an integration of LC-MS metabolomics and constraint-based modeling. Our findings of the effects of CAFs on central carbon metabolic pathways can be summarized as follows (Table 1). First, CAFs induce a glycolytic switch in CRC cells. Compared to normal cells, cancer cells have higher rates of glucose uptake and lactate production even in the presence of oxygen and functional mitochondria, a phenomenon known as the Warburg effect [24, 25]. Comparing CRC cells cultured in CRC media and CAF-conditioned media, we have found that CAFs exacerbate the Warburg effect by enabling cancer cells to uptake glucose and produce lactate at even higher rates. Second, CAFs increase the fluxes through the oxidative arm of the PPP but do not affect the non-oxidative arm of the PPP in CRC cells. The purpose of CAF-induced upregulation of oxidative PPP is perhaps to maintain the redox balance and support the increased demand for nucleotide synthesis in tumor proliferation. Like previous studies, our analysis demonstrates that cells can decouple oxidative and non-oxidative PPP [18, 26]. Third, CAFs inhibit the TCA cycle, increase glutamine uptake, and stimulate glutaminolysis in CRC cells. Our analysis suggests that the enhanced uptake of glutamine in CAF-conditioned media is perhaps not entirely driven by the TCA cycle anaplerosis. At the metabolite level, we have found that CAFs concomitantly switch

**Table 1:** Regulation of CRC metabolism by CAFs. An upward and downward arrows represent a higher or lower median flux, respectively, through the pathway in CRC cells cultured in CAF-conditioned media compared to CRC cells cultured in CRC media.

| Cell | Treatment | Control | Glycolysis | Oxidative PPP | Non-oxidative PPP | TCA Cycle | Glutaminolysis |
| --- | --- | --- | --- | --- | --- | --- | --- |
| KRAS<sup>WT</sup> | CAF-conditioned Media | CRC Media | ↑ | ↑ | - | ↓ | ↑ |
| KRAS<sup>MUT</sup> | CAF-conditioned Media | CRC Media | ↑ | ↑ | ↓ | ↓ | ↑ |

CRC cells to a phenotype characterized by elevated energy production and lactate fermentation. Lactate was traditionally considered a waste product of glycolysis, but an increasing number of studies found that it can play important roles in promoting metastasis, stimulating angiogenesis, inducing immunosuppression, and maintaining the NAD^+^/NADH ratio [27, 28, 29, 30]. Hence, the upregulated conversion of pyruvate to lactate may be driven by the diverse functions of lactate as an oncometabolite [27, 31]. While cancer metabolism is typically associated with high energy demand, our analysis suggests that cancer cells can modulate their biosynthetic pathways in a more complicated manner in order to adapt to their environments.

Our findings are supported by previous studies on tumor-stromal metabolic crosstalk in other cancer types [32, 33]. In agreement with our results, [32] and [33] demonstrated that CAFs enhance extracellular lactate levels in head and neck squamous cell carcinoma (HNSCC) cells via upregulation of glycolysis. Their work further suggests that CAF-secreted HGF activates the glycolytic switch in HNSCC cells via paracrine signaling, whereas HNSCC-cell-secreted basic fibroblast growth factor (bFGF) upregulates oxidative phosphorylation and downregulates glycolysis in CAFs [32, 33]. It will be interesting to validate the roles of HGF and bFGF in CRC-CAF metabolic interactions and identify additional signaling molecules that contribute to CRC metabolic reprogramming via biochemical assays.

By performing single gene deletion analysis, we have also made predictions of knockout metabolic phenotypes. While a large number of *in silico* single-enzyme knockouts result in stronger inhibition of cancer growth in CRC media than in CAF-conditioned media, deletions of HK and G6PD, in particular, result in slower cancer growth in CAF-conditioned media. In the context of metabolic control analysis, this means that HK and G6PD would have higher control coefficients [34]. Biologically, this implies that targeting HK and G6PD suppresses tumor growth by exploiting the metabolic crosstalk between CRC cells and CAFs. Inhibitors of these two enzymes have been explored in the literature. Metformin, a common anti-diabetic drug, was found to inhibit the function of HK in breast and cervical cancer cell lines [35, 36]. Polydatin, a natural molecule found in plants, was demonstrated to be a direct inhibitor of G6PD in HNSCC cell lines [37]. It will be of great interest to validate the efficacy of metformin and polydatin in CRC cell lines and explore the possibility of combining metformin and polydatin with existing CRC therapies.

Method wise, our work demonstrates upFBA as an effective data-driven approach for inferring flux distributions from high-throughput metabolomics data. Metabolite pool size profiling alone may be insufficient for understanding the causes of metabolic alterations due to the intertwined structure of metabolic networks, as we have shown in Section 2.1. While transcriptomics and proteomics data are commonly used to investigate drivers of metabolic/phenotypic differences [38, 39, 40, 41], the expression levels of genes encoding the metabolic enzymes do not fully represent their enzymatic activities. Our method serves to complement the tool box available for metabolism research by making direct use of data obtained at the metabolite level.

We acknowledge some limitations of our study. First, we aim to capture the averaged metabolite dynamics and assume that the rate of change of metabolites is constant between T = 0 hrs and T = 24 hrs. Similar to uFBA [10], more frequent sampling will enable us to capture the fast dynamics more accurately by discretizing metabolite time profiles into time intervals for piecewise simulation. Second, we have only compared CRC media and CAF-conditioned media in this work. Media derived from cocultures of CRC cells and CAFs, however, may diverge from media derived from either cell type alone. It will be interesting to study the metabolism of cancer cells cultured in the media derived from cocultures of CRC cells and CAFs. In addition, upFBA cannot identify the mechanism(s) underlying flux changes, e.g., whether a change in flux is driven by a change in the substrate concentration or a conformational change in the enzyme. Nevertheless, upFBA shows promise as a bridge connecting metabolomics and fluxomics in systems biology and multi-omics studies.

In conclusion, by combining metabolomics data and constraint-based modeling, we have identified metabolic perturbations that exploit the CAF-induced metabolic changes in CRC cells. Our work not only provides quantitative insights that complement experimental studies to develop therapeutic strategies for reducing CRC cell growth but also drives targeted follow-up experiments that can accelerate biological discovery as well as potential clinical translations

## 4. Methods

### 4.1. Cell Culture

#### 4.1.1. Culture Media

Colorectal cancer media (CRC media) are composed of Advanced DMEM/F12 with 10% FBS, 1% penicillin/streptomycin, 1% Glutamax, and 1% HEPES, 100 ng/ml Noggin (Tonbo, 21-7075-U500), 50 ng/ml EGF (Life Technologies, PGH 0313), 10 *µ*M SB202190 (Sigma, 47067), 500 nM A-83-01 (Millipore, 616454-2MG), 10 mM nicotinamide (Sigma, N0636), 1X B-27 (Sigma Aldrich,17504044), 1 mM N-Acetylcysteine (Sigma Aldrich, A7250), and 1X N2 (Sigma Aldrich, 17502048). CRC media conditioned by patient-derived cancer associated fibroblasts (CAF-conditioned media) were made by growing CAFs in 10 cm plates up to 70-80% confluency at standard culture conditions (5% CO_2_, 37 °C). Once the desired confluency was reached, the media were refreshed and allowed to be conditioned by the CAFs for 72 hours. The media were then collected and filtered through a 0.2 *µ*M filter. Conditioned media were aliquoted and stored at −80 °C until used for experiments. CAFs were then collected from the plates and counted with a Bio-Rad T-20 cell counter and trypan blue for normalization purposes. Patient-derived CAFs were isolated from tumor tissue resections of colorectal cancer patients from the USC Norris Comprehensive Cancer Center following Institutional Review Board (IRB) approval and patient consent. The tumor tissue was plated on plastic tissue culture plates to isolate the CAFs and letting the cells grow out over 1-2 passages. These cells were confirmed as CAFs with qPCR and immunofluorescence staining as described previously [42].

#### 4.1.2. Preparation of cells for LC-MS metabolomics

DLD-1 KRAS^WT^ and DLD-1 KRAS^MUT^ cells were obtained from the Yun lab [17] and maintained in McCoy’s 5A media supplemented with 1% penicillin /streptomycin and 10% fetal bovine serum. For LC-MS metabolomics studies, 200,000 cells were seeded in each well of a 6 well plate (3 wells with DLD-1 KRAS^WT^ cells and 3 wells with DLD-1 KRAS^MUT^ cells) in CRC media. Replicates of this plate were prepared for cell counting purposes and for switching the media after 24 hours. After the DLD-1 KRAS^WT^ and KRAS^MUT^ cells were grown for 24 hours in CRC media, intracellular and extracellular metabolite extractions were done as described in Section 4.2 below, switched to CAF-conditioned media, or left in CRC media for an additional 24 hours before metabolite extractions (Figure 1). At each of these time points, plates which were reserved for counting were treated the same way before treated with 0.5% trypsin/EDTA and counted with a TC-20 cell counter.

### 4.2. Mass Spectrometry-Based Metabolomics Analysis

DLD-1 KRAS^WT^ or DLD-1 KRAS^MUT^ cells were seeded in 6-well plates at density of 200,000 cells/well. For extraction of intracellular metabolites, cells were washed on ice with 1 ml ice-cold 150 mM ammonium acetate (NH_4_AcO, pH 7.3). 1 ml of −80 °C cold 80% MeOH was added to the wells, and samples were incubated at −80 °C for 20 minutes. Then cells were scraped off, and supernatants were transferred into microfuge tubes. Samples were pelleted at 4 °C for 5 minutes at 13k rpm. The supernatants were transferred into LoBind Eppendorf microfuge tube; the cell pellets were re-extracted with 200 *µ*l ice-cold 80% MeOH, spun down and the supernatants were combined. Metabolites were dried at room temperature under vacuum and re-suspended in water for LC-MS run. For extraction of extracellular metabolites, 20 *µ*l of cell-free blank and conditioned media samples were collected from wells. Metabolites were extracted by adding 500 *µ*l −80 °C cold 80% MeOH, dried at room temperature under vacuum and re-suspended in water for LC-MS analysis.

For each experiment, biological triplicate samples were randomized and analyzed on a Q Exactive Plus hybrid quadrupole-Orbitrap mass spectrometer coupled to an UltiMate 3000 UHPLC system (Thermo Scientific). The mass spectrometer was run in polarity switching mode (+3.00 kV/-2.25 kV) with an m/z window ranging from 65 to 975. Mobile phase A was 5 mM NH_4_AcO, pH 9.9, and mobile phase B was acetonitrile. Metabolites were separated on a Luna 3 *µ*m NH2 100 Å (150 × 2.0 mm) column (Phenomenex). The flowrate was 300 *µ*l/min, and the gradient was from 15% A to 95% A in 18 minutes, followed by an isocratic step for 9 minutes and re-equilibration for 7 minutes.

Metabolites were detected and quantified as area under the curve based on retention time and accurate mass (≤ 5 ppm) using the TraceFinder 3.3 (Thermo Scientific) software. Extracellular data were normalized to integrated cell number, which was calculated based on cell counts at the start and end of the time course and an exponential growth equation. Intracellular data was normalized to the cell numbers at the time of extraction. Additionally, both intracellular and extracellular metabolomics data were median-normalized. We calculated the fold changes of metabolite levels by dividing the final abundance of the metabolite (denoted as the T = 24 hrs timepoint) by its baseline condition in the CRC media (denoted as the T = 0 hrs timepoint). Differential abundance analysis was implemented using the LIMMA Bioconductor package (R version 3.6.2) [13].

### 4.3. Constraint-Based Modeling

To infer flux distributions, we performed upFBA [10, 11]. Compared to existing flux-inference methods, upFBA is more suitable for our study for several reasons. First, upFBA does not require detailed knowledge of enzymatic kinetics. In a typical LC-MS metabolomics experiment, only a subset of metabolites can be experimentally identified and quantified. However, the number of parameters to fit in ODE-based kinetic models are likely to far exceed the number of metabolites for which data are available [43, 44, 45, 46]. Second, upFBA can be used to examine media-induced metabolic alterations. As the primary goal of our study is to understand the effects of CAF-conditioned media on CRC metabolism, manipulating the media would defeat the purpose of our study. Hence, ^13^C metabolic flux analysis (^13^C-MFA) is not particularly useful here because a ^13^C-labeled substrate needs to be introduced to the culture medium in a tracing experiment [47, 48, 49]. Third, upFBA does not require the steady-state assumption. Both steady-state FBA and ^13^C-MFA assume that metabolites in the system are at steady state, so these analyses are difficult to interpret when cells undergo a response to perturbations such as a media switch [50]. Fourth, unlike uFBA [10], upFBA does not require absolute quantification of metabolites. Instead, it combines knowledge of initial metabolite concentrations and measurements of fold changes to estimate mass balance constraints. Finally, upFBA increases the reliability of the estimated flux distributions by repeatedly sampling the initial metabolite concentrations and generating mass balance constraints from the sampled concentrations and measured fold changes of metabolites.

Our upFBA approach stems from uFBA developed by [10] but differs in the specification of mass balance constraints as well as objective functions. We specified mass balance constraints by integrating the estimates of initial metabolite concentrations from literature and measurements of relative metabolite changes from our experimental data. Our approach operates in following steps (Figure 8). First, we sampled initial metabolite concentrations (T = 0 hrs) 100 times from the range estimates reported in the literature. Due to experimental limitations, not every intermediate can be measured. Experimental data are available for 14 metabolites in central metabolism, and the range estimates of initial concentrations are available for 7 out of the 14 metabolites in cancer cells: glucose 6-phosphate (G6P), fructose 1,6-bisphosphate (FBP), glyceraldehyde 3-phosphate (G3P), phosphoenolpyruvate (PEP), lactate, glutamine, and glutamate [51, 52]. (See Table S1 for range estimates). Let *l*_*i*_ and *u*_*i*_ denote the lower and upper bounds of the initial concentration of the *i*-th metabolite for which estimates are available (*i* = 1, 2, …, 7). The initial concentration of the *i*-th metabolite in the *j*-th sampling (*j* = 1, 2, …, 100), 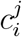, needs to stay between the lower and upper bounds, i.e., 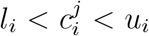. Thus, the initial concentrations of all metabolites for which range estimates are available in the *j*-th sampling, 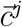, satisfy

**Figure 8:**
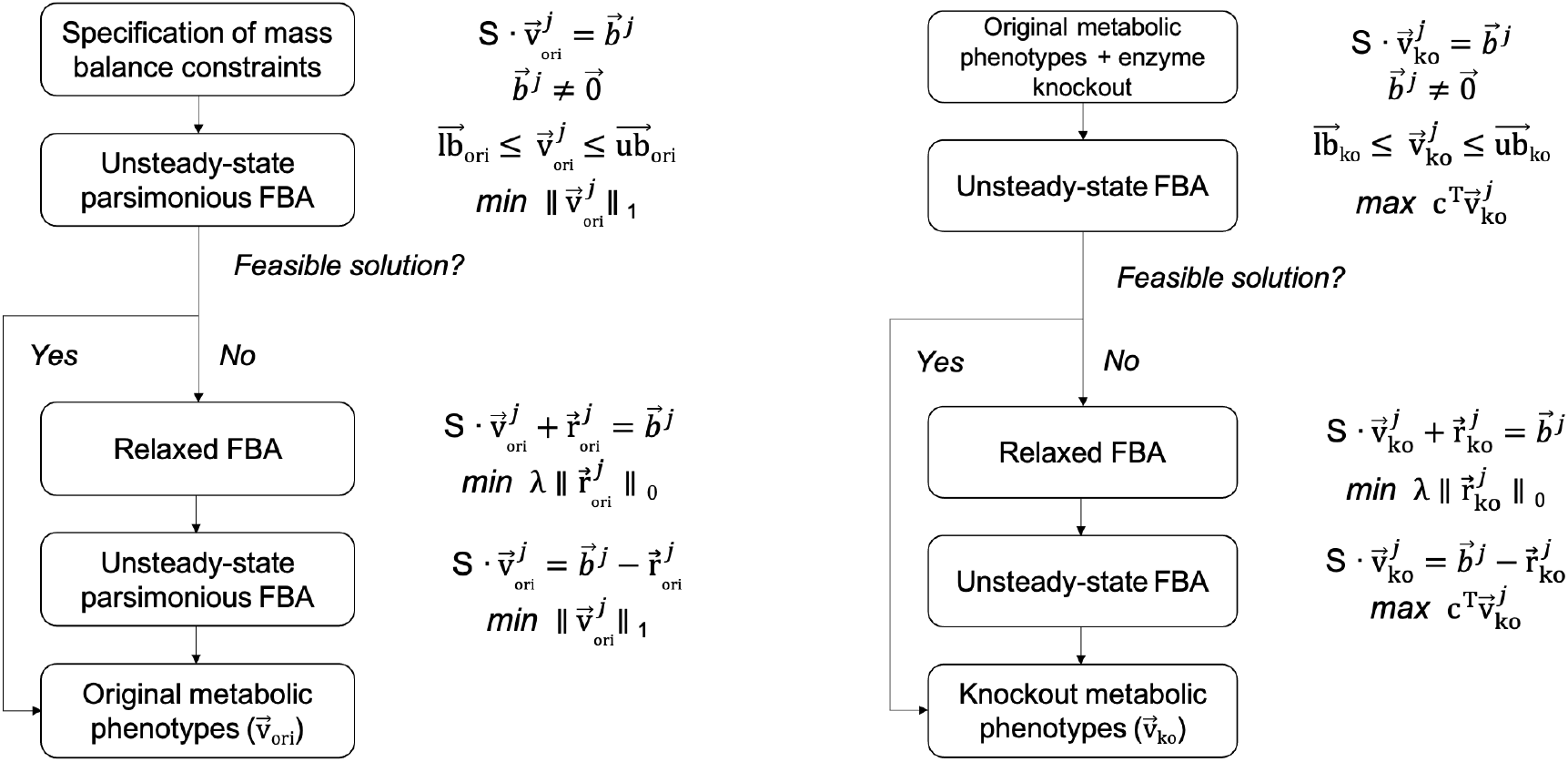
A constraint-based modeling approach to predicting metabolic phenotypes. The same workflows are carried out repeatedly for each set of initial metabolite concentration that was sampled (*j* = 1, 2, …100).

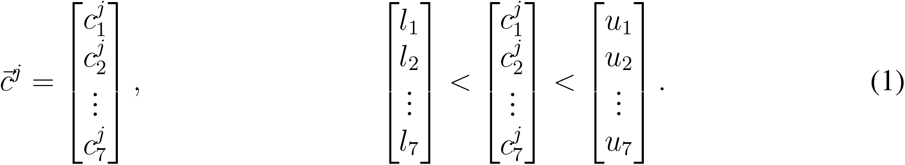

Let *S*_*i,T* =0_ and *S*_*i,T* =24_ denote the normalized abundances of the *i*-th metabolite at T = 0 hrs and T = 24 hrs. The fold change of the *i*-th metabolite is defined as the ratio of *S*_*i,T* =24_ to *S*_*i,T* =0_. The mass balance constraint on the *i*-th metabolite in the *j*-th sampling, which represents the absolute change in abundance between T = 0 hrs and T = 24 hrs, can be expressed as

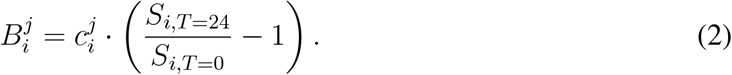

Thus, the mass balance constraints on all metabolites for which range estimates are available in the *j*-th sampling, 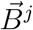, are specified as

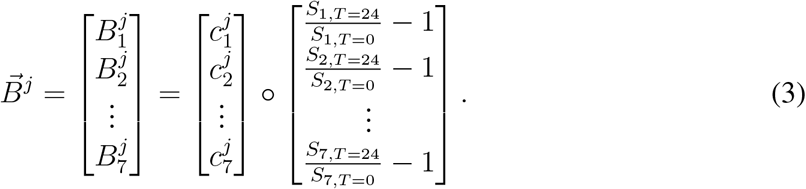

Next, we used linear programming to calculate a solution that satisfies the mass balance constraints and flux constraints. To avoid increasing the degrees of freedom of the model, we began by relaxing steady-state conditions for the seven metabolites for which range estimates are available and assuming steady-state conditions for the rest [10] (Figure 8). In other words, the mass balance constraints on all metabolites in our model in the *j*-th sampling, 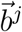 (Figure 8), can be expressed as

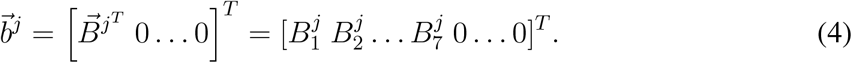

We also applied flux constraints. The uptake/secretion rates of glucose, lactate, and glutamine were experimentally measured, so the fluxes through GLUT, MCT, and ASCT2-mediated transport were all set to fixed values. (See Table S2 for data). The units of uptake/secretion rates were converted from mmol/(cell·hr) to mM/hr under the assumption that each DLD-1 cell has an approximate volume of 1000 *µ*M^3^ [53]. Biomass growth rates were calculated based on integrated cell counts measured over three days and also set to fixed values in our models. The upper bounds on all remaining fluxes were set to 500 mM/hr. The lower bounds on the fluxes through the remaining reversible and irreversible reactions were set to −500 mM/hr and 0 mM/hr, respectively. A flux of 500 mM/hr was chosen to represent an infinite flux, as such a value is too high to achieve in a real biological system. Lastly, a flux split between glycolysis and the PPP was constrained to be 90*/*10 based on a previous ^13^C tracing study performed on cancer cell lines [54].

Unlike uFBA, which maximizes the biomass objective function, upFBA minimizes the total sum of fluxes through the metabolic network in the *j*-th sampling, 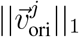 (Figure 8) [11]. Based on the optimality principle, it is reasonable to assume that cells seek to maintain a pre-specified growth rate at the lowest metabolic cost possible [11]. In cases where models failed to yield feasible solutions, we performed relaxed FBA (rFBA) which minimizes the number of unmeasured metabolites for which mass balance constraints need to be relaxed [55] (Figure 8). As suggested by [10], implementing this objective to minimize the sum of the fluxes through the metabolic network produces more accurate results than other common objective functions used in this context. The mass balance constraints for just three metabolites needed to be relaxed in the various upFBA runs (glucose [Glc], fructose 6-phosphate [F6P], and sedoheptulose-7-phosphate [S7P]), and the resulting number of times mass balance constraints were relaxed on these metabolites is shown in Figure S4. Following rFBA, we performed another round of upFBA with the relaxed constraints to estimate the reaction fluxes (Figure 8). Scripts relevant to upFBA, including the reaction network we built for central carbon metabolism, can be found on GitHub.

The single gene deletion analysis operates similarly. We simulated gene deletion by constraining the deleted reaction to zero between T = 0 hrs and T = 24 hrs. Mathematically, this means that the upper bound and lower bound for the deleted reaction in the *j*-th sampling were set to 0 mM/hr. In contrast, mass balance constraints, 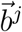 and the remaining flux constraints remained the same, except the constraint on the biomass growth rate (Figure 8). uFBA was then performed to predict the maximal biomass growth rate that could be achieved under the new flux constraints (Figure 8). Similar to the steps outlined above, we utilized rFBA in cases where models did not produce feasible solutions [55], following which we performed another round of uFBA to predict the maximal biomass growth rate possible (Figure 8). Scripts relevant to single gene deletion analysis can be found on GitHub.

Both upFBA and uFBA were implemented in MATLAB R2019b using the COBRA Toolbox v3.0 [55, 56]. Linear programming was solved using the GLPK solver [57]. The resulting fluxes from upFBA and biomass growth rates from uFBA were compared between cells cultured in CRC media and CAF-conditioned media via the Wilcoxon rank sum test [16], which either accepts or rejects the null hypothesis that a reaction has an equal median flux under two media conditions. The adjusted p-values were subsequently computed from the p-values using the Bonferroni correction.

## Supplementary Information

Supplementary tables, figures, and files can be found online together with this article. MATLAB scripts are available on GitHub: https://github.com/FinleyLabUSC/CRC-Cell-Constraint-Based-Metabolic-Model.

## Acknowledgements

This work was supported by the NIH National Cancer Institute grant 1U01CA232137.

MATLAB scripts are available on GitHub: https://github.com/FinleyLabUSC/CRC-Cell-Constraint-Based-Metabolic-Model

