## Supplementary Information for "Elucidating Tumor-stromal Metabolic Crosstalk in Colorectal Cancer through Integration of Constraint-Based Models and LC-MS Metabolomics"

SUPPLEMENTARY TABLES AND FIGURES

| Metabolite | Lower (mM) | Upper (mM) | Ref. |
|---|---|---|---|
| G6P[c] | 1.68 | 6 | [2] |
| FBP[c] | 0.16 | 1.08 | [2] |
| G3P[c] | 0.24 | 0.84 | [2] |
| PEP[c] | 0.24 | 0.84 | [2] |
| lactate[c] | 6.4 | 19.2 | [2] |
| Gln[i] | 0.56 | 1.08 | [1] |
| Glu[m] | 0.88 | 1.56 | [1] |

**Table S1:** Bounds for initial metabolite concentrations of CRC cells, including both $KRAS^{WT}$ and $KRAS^{MUT}$ CRC cells used in upFBA. The concentration ranges of CRC cells were specified based on published measurements obtained from various cell lines, including breast cancer cell extracts [1] and human cervical cancer cells [2]. To account for additional uncertainty, we increased and decreased the upper and lower bounds reported in the literature, respectively, by 20%.

| | Glucose ($mM{\cdot}hr^{-1}$) | Lactate ($mM{\cdot}hr^{-1}$) | Glutamine ($mM{\cdot}hr^{-1}$) | Biomass ($hr^{-1}$) |
|---|---|---|---|---|
| $KRAS^{WT}$ (CRC media) | 0.223 (0.038) | 0.283 (0.016) | 0.003 (0.002) | 0.035 (0.001) |
| $KRAS^{MUT}$ (CRC media) | 0.210 (0.024) | 0.234 (0.004) | 0.003 (0.001) | 0.034 (0.003) |
| $KRAS^{WT}$ (CAF-cond. media) | 0.573 (0.087) | 0.784 (0.041) | 0.052 (0.025) | 0.033 (0.001) |
| $KRAS^{MUT}$ (CAF-cond. media) | 0.927 (0.135) | 1.117 (0.144) | 0.105 (0.049) | 0.033 (0.003) |

**Table S2:** Measurements of glucose uptake rates, lactate secretion rates, glutamine uptake rates, and cell proliferation rates. Glucose uptake, lactate secretion, and glutamine uptake rates between T = 0 hrs and T = 24 hrs were measured via LC-MS metabolomics. Biomass growth rates were measured via cell proliferation assays. These values were supplied to our upFBA models as constraints. Glucose uptake, lactate secretion, and glutamine uptake rates have units of $mM{\cdot}hr^{-1}$. Biomass growth rates have units of $hr^{-1}$. Numbers inside parentheses stand for standard deviations.

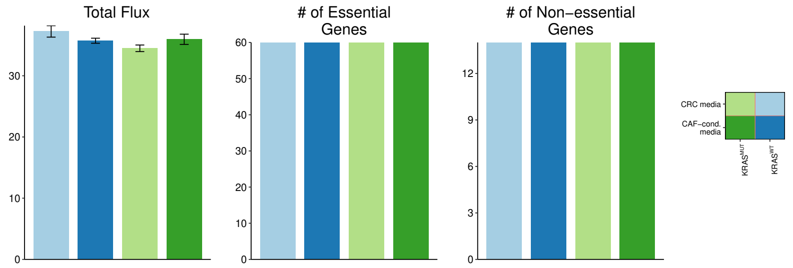


**Figure S1:** Minimized sum of fluxes and numbers of essential (non-essential) reactions. Total fluxes are represented by mean +/- standard deviation. Essential (non-essential) reactions are defined as reactions that (do not) maintain a non-zero flux under at least one set of mass balance constraints.

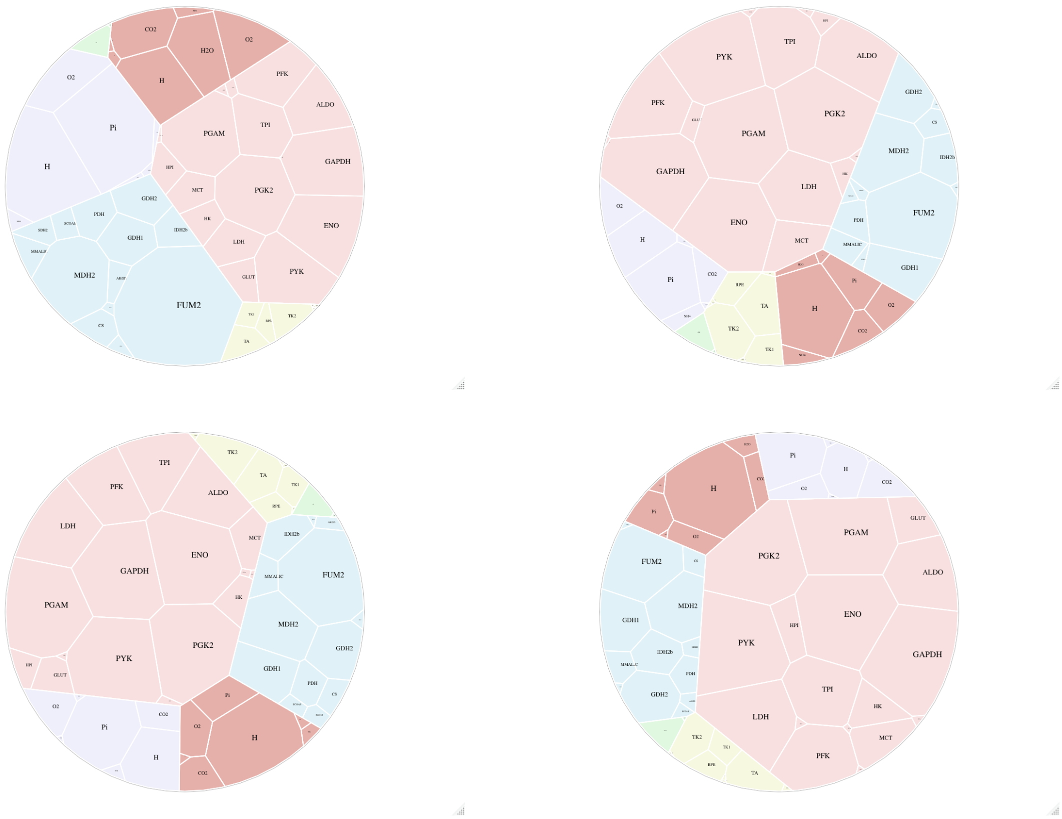


**Figure S2:** Voronoi treemaps showing the flux distributions of KRAS$^{WT}$ CRC cells cultured in CRC media (upper left), KRAS$^{MUT}$ CRC cells cultured in CRC media (upper right), KRAS$^{WT}$ CRC cells cultured in CAF-conditioned media (bottom left), and KRAS$^{MUT}$ CRC cells cultured in CAF-conditioned media (bottom right). Tile size represents the median absolute flux value of a reaction, i.e., a larger tile corresponds to a higher median flux. Tile color represents the pathway. Color code: pink, glycolysis; yellow, PPP; blue, TCA cycle; green, glutaminolysis; purple, nuclear transport; red, cell membrane transport.

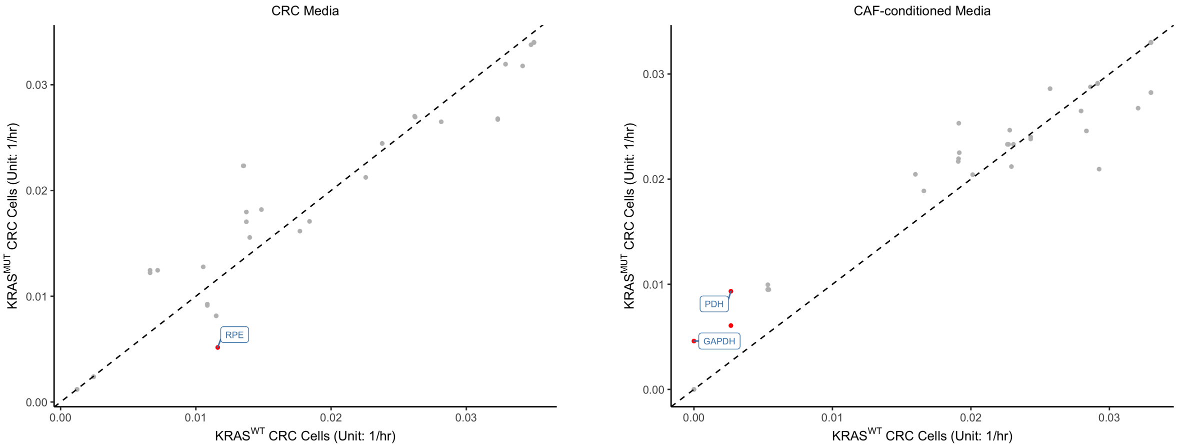


**Figure S3:** Maximal biomass growth rates of CRC cells cultured in CRC media and CAF-conditioned media upon gene deletions predicted by uFBA. The *x*- and *y*- coordinates of each point represent the median maximal biomass growth rates of $KRAS^{WT}$ and $KRAS^{MUT}$ CRC cells, respectively when the same enzyme is inhibited. The dashed diagonal line represents equal median rates in both media. Gene deletions resulting in significantly different biomass growth rates between $KRAS^{WT}$ and $KRAS^{MUT}$ CRC cells are highlighted in red (adjusted p-value $< 0.001$, fold change $> 2$). Enzymes belonging to glycolysis, PPP, the TCA cycle, and glutaminolysis are labeled.

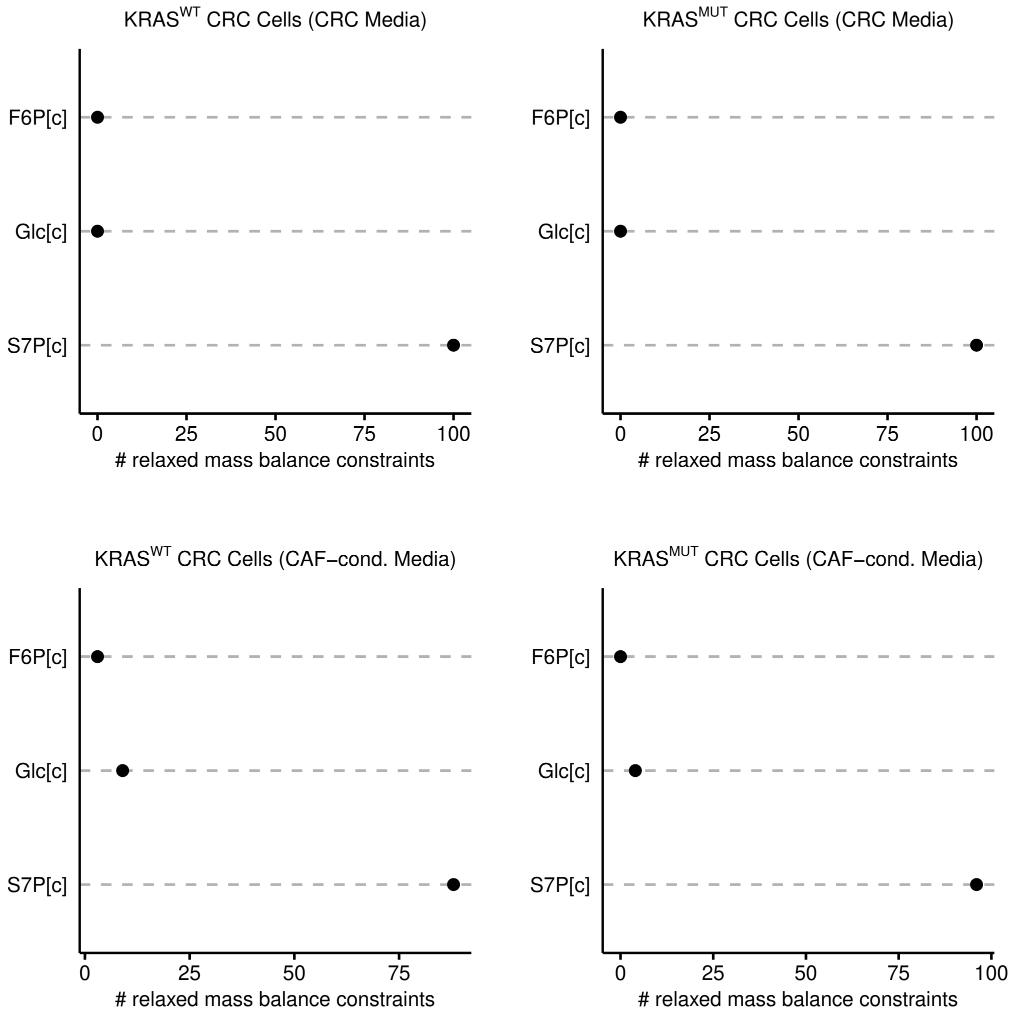


**Figure S4:** Number of times mass balance constraints were relaxed on unmeasured metabolites out of 100 models for KRAS$^{WT}$ and KRAS$^{MUT}$ CRC cells cultured in CRC media and CAF-conditioned media. For metabolites not in the plots, no deviation from steady states was found under any set of mass balance constraints.
